# SMCHD1 activates the expression of genes required for the expansion of human myoblasts

**DOI:** 10.1101/2024.02.04.578809

**Authors:** Matthew Man-Kin Wong, Ed Gardner, Lynn A. Megeney, Charles P. Emerson, Davide Gabellini, F. Jeffrey Dilworth

## Abstract

SMCHD1 is an epigenetic regulatory protein known to modulate the targeted repression of large chromatin domains. Diminished SMCHD1 function in muscle fibers causes Facioscapulohumeral Muscular Dystrophy (FSHD2) through derepression of the *D4Z4* chromatin domain, an event which permits the aberrant expression of the disease-causing gene *DUX4*. Given that SMCHD1 plays a broader role in establishing the cellular epigenome, we examined whether loss of SMCHD1 function might affect muscle homeostasis through additional mechanisms. Here we show that acute depletion of SMCHD1 results in a DUX4-independent defect in myoblast proliferation. Genomic and transcriptomic experiments determined that SMCHD1 associates with enhancers of genes controlling cell cycle to activate their expression. Amongst these cell cycle regulatory genes, we identified LAP2 as a key target of SMCHD1 required for the expansion of myoblasts, where the ectopic expression of LAP2 rescues the proliferation defect of SMCHD1-depleted cells. Thus, the epigenetic regulator SMCHD1 can play the role of a transcriptional co-activator for maintaining the expression of genes required for muscle progenitor expansion. This DUX4-independent role for SMCHD1 in myoblasts suggests that the pathology of FSHD2 may be a consequence of defective muscle regeneration in addition to the muscle wasting caused by spurious DUX4 expression.

Chromatin modifier SMCHD1 regulates the proliferation of human myoblasts

SMCHD1 binds to enhancers of cell cycle regulatory genes to activate their expression

Lap2 is an SMCHD1 direct target gene that regulates the expansion of human myoblasts.

## INTRODUCTION

The *Structural maintenance of chromosomes flexible hinge domain containing 1* (*SMCHD1*) gene encodes an epigenetic modifier first identified in a screen for genes involved in transgene array silencing (1). SMCHD1 is best known for its role in X chromosome inactivation (XCI) where it plays an essential role in merging the intermediate compartments to ensure proper gene silencing (2, 3). SMCHD1 contributes to the establishment of heterochromatin by regulating DNA methylation, where it is required for Dnmt3b-mediated methylation of CpG islands on the inactive X chromosome (4), while also inhibiting the activity of Tet DNA demethylases (5). SMCHD1 can also actively repress gene transcription by limiting long-range chromatin interactions where it acts to deplete CTCF and cohesin from the topologically associated domain (TAD) boundaries at specific loci, including the Hox and protocadherin gene clusters (3, 6–8).

Loss of function mutations in the *SMCHD1* gene are observed in a subgroup of patients with the muscle-wasting condition Facioscapulohumeral Muscular Dystrophy (FSHD), one of the most frequent neuromuscular diseases. However, the predominant cause of FSHD is a deletion of the subtelomeric *D4Z4* tandem DNA array on chromosome 4. In FSHD1 this contraction of the *D4Z4* repeat to less than 11 copies induces an epigenetic de-repression of the locus, which results in the ectopic expression of an RNA encoding double homeobox 4 (DUX4), a transcription factor whose presence in muscle fibers induces apoptosis (9–11). Based on this mechanism of action, it was later demonstrated that the reduced SMCHD1 function in muscle-induced de-repression of the *D4Z4* tandem array to permit the expression of DUX4 from patients having a relatively short FSHD locus (up to 16 copies of *D4Z4*) which leads to the onset of FSHD2 (12–14). Besides epigenetic de-repression of the D4Z4 locus, another prerequisite for the onset of FSHD is the presence of 4qA alleles with a polyA signal downstream of the last copy of the *D4Z4* which permit the expression of stable *DUX4* mRNA (15–16), while contractions of a 4qB *D4Z4* locus are non-permissive for FSHD. The facts that FSHD2 patients still have substantially shorter *D4Z4* locus than that of the unaffected population and that FSHD-permissive 4qA alleles are required for the onset of FSHD have provided strong evidence that SMCHD1 mutations and shortened *D4Z4* array length both contribute to the de-repression of the *D4Z4* locus, and spurious activation of the DUX4 transcription factor leads to muscle wasting in FSHD. Nevertheless, patients who have mutations of *SMCHD1* in addition to a contraction of the *D4Z4* locus often display a more severe muscle-wasting phenotype (17–18). This raises the question of the potential for the general epigenetic silencer SMCHD1 to have additional roles in muscle cells that might exasperate the disease phenotype.

Here we set out to investigate the function of SMCHD1 in human myoblasts that might affect the health of muscle independently of DUX4 expression. For this, we took advantage of the fact the susceptibility of individuals to FSHD depends upon a specific polymorphism within the *D4Z4* locus since the 4qA chromosome variant creates a polyadenylation signal which is necessary to stabilize *DUX4* transcripts (16, 19). We thus depleted SMCHD1 in cells derived from a healthy individual with a 4qB polymorphism at the *D4Z4* locus on both chromosomes that cannot give rise to stable *DUX4* transcripts (20). Using this approach, we uncovered a previously unappreciated DUX4-independent role for SMCHD1 as a co-activator of genes involved in cell cycle progression during myogenic progenitor cell expansion.

## MATERIAL AND METHODS

### Reagents and software

The names, sources, and catalogue numbers of the reagents used in the experiments are listed in Supplementary Table S1. All the software and databases used in the data analyses are listed in Supplementary Table S2.

### Cell culture

Immortalized non-FSHD myoblast cell line WS234 (donor 15Vbic) (20) was obtained from the Wellstone Muscular Dystrophy Cooperative Research Center for FSHD, University of Massachusetts Medical School, USA. This cell line was previously tested to possess a 4qB D4Z4 chromosome variant on both chromosomes and have no detectable *DUX4* mRNA expression (21). In proliferating condition, cells were cultured in LHCN medium [4:1 DMEM : Medium 199 supplemented with 15% BGS (v/v), 0.03 ug/ml Zinc Sulfate, 1.4 ug/ml vitamin B12, 0.055 ug/ml dexamethasone, 2.5 ng/ml HGF, 10 ng/ml FGF, 0.6X penicillin/streptomycin, 20 mM HEPES (pH 7.4)] on plates coated with 0.1% gelatin (w/v) coated plates at 37 ° C in 5% CO2 incubator. HEK293T cells were cultured in DMEM growth medium [DMEM supplemented with 10% BGS and 1x penicillin/streptomycin], at 37°C in 5% CO2 incubator.

### Plasmid Generation

The plasmid vector pDONR223 containing the open reading frame of human *LAP2β* was purchased from the genome editing and molecular biology facility of the University of Ottawa. The open reading frame of *LAP2β* was amplified by PCR using the forward primer 5’- catggaggatccgccaccatgccggagttcctggaag-3’ and the reverse primer 5’- cgtatggtacctcattagttggattttctagggtca-3’. The resulting PCR product was cloned in the lentiviral vector pLenti-EF1a-Blank and the sequence of *LAP2β* was confirmed by Sanger sequencing.

### Preparation of lentivirus

HEK293T cells were transfected with the pMD2.G, psPAX2, and shRNA vector plasmids (Supplementary Table S3) using polyethylenimine as previously described (22). Lentivirus-containing medium supernatant was centrifuged at 126, 086 x g for 2.5 hours at 4°C using Optima L-100 XP ultracentrifuge (Beckman Coulter), and the pellet was resuspended in LHCN medium. To transduce the cells, purified lentivirus and 8 ug/ml of polybrene was added to the culture medium. The cells were incubated for 16 hours at 37°C and then fresh LHCN medium was replaced.

### Growth curve, EdU proliferation essay

The cell proliferation rate was determined by EdU incorporation in newly synthesized DNA as well as cell counting. Three days after lentiviral transduction, an equivalent number of cells were seeded on each well of a gelatin-coated 35 mm plate. Fresh LHCN medium was replaced every 2 days. Five days after lentiviral transduction, 10 µM EdU was added to the culture medium for 90 minutes at 37°C. Cells were fixed to perform immunostaining according to the manufacturer’s protocol of the EdU staining kit. For cell counting experiments, cells were trypsinized and resuspended LHCN medium. Trypan blue solution was added and the mixture was loaded to a hemocytometer for cell counting. The cell number from each well was calculated from the average of 6 independent hemocytometer counting.

### RNA-Seq and data analysis

Five days after *SMCHD1* or non-silencing lentiviral transduction, RNA was collected using a PureLink RNA mini kit according to the manufacturer’s protocol. The depletion of rRNA was performed using the Ribo-Zero Magnetic Gold Kit, and the RNA library was prepared using KAPA stranded RNA-Seq kit according to the manufacturer’s protocol. Sequencing was performed on an Illumina HiSeq4000 platform. RNA-Seq reads from fastq files were aligned to human hg38 genome using the STAR, counts were summarized using featureCounts function in Rsubread and used as input for DESeq2 to identify the differentially expressed genes. RNA-Seq datasets downloaded from GEO were analysed using the same settings as SMCHD1 depletion RNA-Seq dataset. Volcano plots were made using EnhancedVolcano, dot plots were made using ggplot2 and heatmaps using pheatmap in R. Gene ontology analysis was performed using DAVID, and values were used as inputs for dot plots.

### Gene Set Enrichment Analysis

The down-regulated or up-regulated genes after SMCHD1 knockdown (adjusted p <= 0.05) were used as two separate gene set for Gene Set Enrichment Analysis (GSEA). The analysis was performed using the GSEA software according to the user guide with the number of permutations set to 1000.

### Binding and Expression Target Analysis (BETA)

The analysis was done following the user manual. Peaks within 150 kbp of TSS and differentially expressed genes with adjusted p value <= 0.05 were selected for the analysis, without using CTCF as a boundary to filter peaks around genes.

### ChIP-Seq and Data Analysis

The SMCHD1 ChIP-Seq was performed as previously described (23) with some modifications. Briefly, proliferating WS234 myoblasts were fixed in 1% (w/v) formaldehyde for 20 minutes at room temperature and quenched with 125 mM glycine for 10 minutes at room temperature. Cells were rinsed with PBS, collected using cell scrapers, centrifuged, and resuspended in swelling buffer [50 mM Tris base (pH 8.0), 300 mM sucrose, 10 mM NaCl, 0.5% NP-40 (v/v)]. Cells were incubated for 30 minutes in swelling buffer at 4°C, and sonicated using Bioruptor (Diagenode) for 2 cycles (HI power, cycles of 30 sec on / 1 min off). Nuclei were spun down at 1, 000 xg for 5 minutes at 4°C, resuspended in sonication buffer [50 mM HEPES (pH 7.9), 140 mM NaCl, 1 mM EDTA, 1% Triton X-100 (v/v), 0.1% Na-deoxycholate (w/v), 1% SDS (w/v)], and sonicated for 13 cycles using the same settings as the previous step. Next, 7.5 ug of SMCHD1 antibody or normal rabbit IgG was added to 250 ug of precleared chromatin and incubated for 3 hours at room temperature. The chromatin was then washed and purified as described previously (23). Library was prepared using Kapa Hyperprep kit and paired-end sequencing was performed on an Illumina HiSeq4000 platform. The reads were aligned to the human hg38 genome using bowtie2 and peaks were called by MACS2 using a q value cutoff = 0.05. Meta profile plots of ChIP-Seq signal around SMCHD1 peaks were generated using EnrichedHeatmap and ggplot2 in R. Raw ChIP-Seq fastq files downloaded from GEO were analysed using the same settings.

### Motif identification using HOMER (Hypergeometric Optimization of Motif EnRichment)

The bed files generated by MACS2 were converted to fasta files using “getfasta” function in BEDTools. The converted fasta files were used as input files for HOMER analysis using the “findMotifs.pl” function.

### Statistical Analyses

For RNA-Seq, differentially expressed genes were defined as genes with a fold change > 0.5 and an adjusted p-value <= 0.05, unless otherwise specified. For the identification of the ChIP-Seq peak identification using MACS2, the cutoff value q value cutoff = 0.05. For overlapping ChIP-Seq peaks between two datasets, p values and Z scores were calculated by permutation test using regioneR with number of permutations = 1000. Two-way ANOVA (for cell counting experiments) and student’s two-tailed t-test (for other experiments) were performed using the statistical graphing software GraphPad Prism v6.0, and *p<0.05, **p<0.01, ***p<0.001, ****p<0.0001, ns = not significant.

## RESULTS

### SMCHD1 is required for efficient proliferation of human myoblasts

To determine the roles of SMCHD1 that are independent of DUX4 in adult myogenesis, we performed an acute depletion of SMCHD1 in immortalized human WS234 myoblasts (20). WS234 cells were chosen as they possess the non-pathogenic 4qB *D4Z4* allele on both copies of chromosome 4, preventing the expression of a stable *DUX4* mRNA transcript (21). We opted to use an shRNA-mediated approach for depletion to prevent the selection of clones that had adapted to the complete loss of SMCHD1. Using three different shRNAs targeting distinct regions of the *SMCHD1* transcript, we observed a reduction in mRNA and protein accumulation that varied between 50 and 90% when examined 5 days after lentivirus infection (Supplementary Figure S1A and S1B). These reduced levels of SMCHD1 would be consistent with the haploinsufficiency in FSHD patients possessing a heterozygous mutation of the gene. Phenotypically, the reduction in SMCHD1 levels in cells caused the muscle progenitor cells to adopt a less elongated morphology compared to the controls (Figure 1A). This change in cell appearance suggests that SMCHD1 may be important for myoblast function.

**Figure 1.**
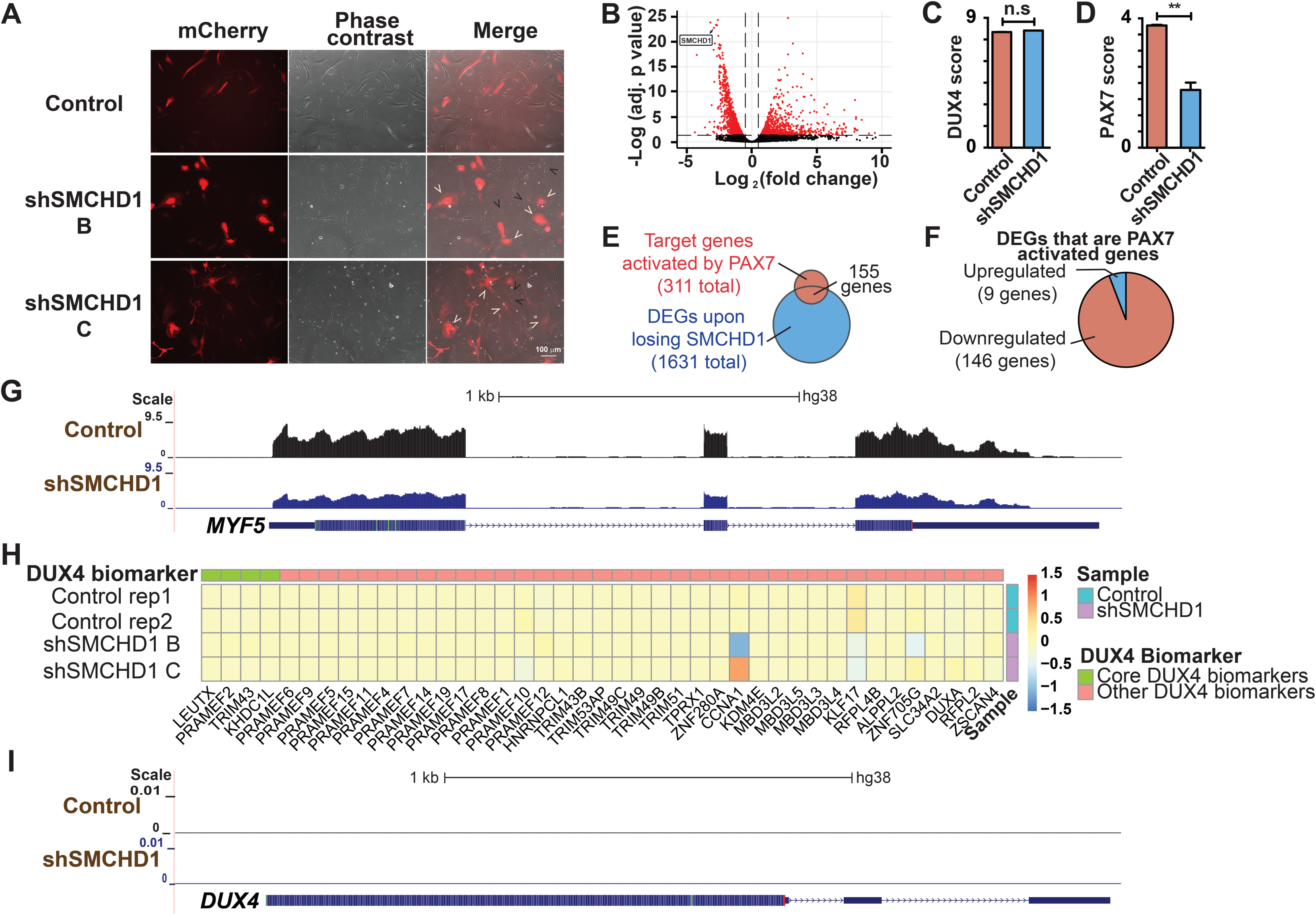
Loss of SMCHD1 leads to FSHD-like repression of PAX7 target genes in myoblasts without DUX4 target genes activation. **(A)** Nine days after lentiviral transduction, cells were observed under fluorescence microscope. (White arrows) Transduced mCherry^+^ *SMCHD1* shRNA expressing cells. (Black arrows) Cells that do not express shRNA expression. **(B)** Volcano plot showing the DEGs after SMCHD1 knockdown. Genes with adjusted p value <= 0.05 and log2 fold change > 0.5 were labelled red. **(C)** DUX4 score of samples transduced with non-silencing scrambled shRNA (Control) or *SMCHD1* targeting shRNA (shSMCHD1) based on DUX4 targets identified by Choi *et al*. 2016 (25). **(D)** PAX7 scores of samples transduced with non-silencing scrambled shRNA (Control) or *SMCHD1* targeting shRNA (shSMCHD1). **(E)** Venn diagram of the target genes activated by PAX7 and differentially expressed upon SMCHD1 depletion. **(F)** Number of PAX7 activated genes activated by PAX7 that were down-regulated (red) and up-regulated (blue) upon SMCHD1 depletion. **(G)** UCSC genome browser tracts showing RNA-Seq reads of samples transduced with non-silencing scrambled shRNA (Control) or *SMCHD1* targeting shRNA (shSMCHD1) near the *MYF5* gene. **(H)** Heatmap showing the expression of DUX4 target genes identified by Yao et. al. (27) upon SMCHD1 depletion. **(I)** UCSC genome browser tracts showing RNA-Seq reads of samples transduced with non-silencing scrambled shRNA (Control) or *SMCHD1* targeting shRNA (shSMCHD1) near the *DUX4* gene.

### Loss of SMCHD1 leads to aberrant expression of PAX7 target genes in myoblasts

To understand how the loss of SMCHD1 impairs myoblast function, we performed RNA-sequencing analysis on myoblasts depleted of SMCHD1 using two distinct shRNAs. Analysis of gene expression changes in myoblasts depleted of SMCHD1 identified 1631 differentially expressed genes (932 down-regulated and 699 up-regulated) (Figure 1B and Supplementary File S1). Though we were using myoblasts which possess only the 4qB *D4Z4* allele, we wanted to confirm the absence of DUX4 in our myoblasts. Indeed, no transcripts of *DUX4* were detected in our deep-sequencing of RNA from WS234 cells under any conditions tested. Due to the difficulty of detecting *DUX4* transcripts in muscle cells, measurement of DUX4-target gene expression is often used as a surrogate for detecting the presence of DUX4 (24). Measuring the previously described “DUX4 score” (24), we observed that SMCHD1 knockdown did not significantly change the expression of previously established DUX4 biomarkers (25–27) (Figure 1C and Supplementary Figure S1C). Reasoning that the loss of SMCHD1 may only change the expression of the most robust DUX4-FSHD biomarkers, we re-assessed a DUX4 score using only the top biomarkers identified by Yao et al. (27). Again we found that loss of SMCHD1 did not significantly change the expression of the four core sets of DUX4-FSHD biomarkers (Figure 1H). Thus, the proliferation defects observed in the absence of SMCHD1 are not the result of induced DUX4 expression.

A hallmark of myoblasts from FSHD patients is the repression of Pax7 target genes (28). As such, we next examined whether the expression of known Pax7 target genes was altered upon depletion of SMCHD1. For this purpose, we calculated a PAX7 score (24) based on changes in transcript abundance upon SMCHD1 depletion (Figure 1D). We observed that ∼50% of the genes normally activated by Pax7 in myoblasts show reduced expression in myoblasts depleted of SMCHD1 (Figure 1E-G), while ∼11% of genes repressed by PAX7 expression show increased expression upon SMCHD1 depletion (Supplementary Figure S1D-E). Therefore, our data showed that SMCHD1 depletion reproduces some of the altered gene expression as observed in myoblasts-derived from FSHD patients despite the fact that DUX4 itself is not expressed (Figure 1I).

To better understand how the loss of SMCHD1 was affecting myoblast function, we performed gene ontology (GO) enrichment analysis. Consistent with results observed in FSHD patients with *SMCHD1* mutations (29), loss of the epigenetic modifier in myoblasts results in the upregulation of genes associated with the extracellular matrix (Figure 2A and Supplementary Figure S2A). Though SMCHD1 is known as an epigenetic silencer, we were intrigued by the large number of genes involved in cell cycle (DNA replication, nucleosome assembly and G1/S transition) that were down-regulated upon loss of SMCHD1 (Figure 2B). Using RT-qPCR, we confirmed that SMCHD1-depletion leads to the downregulation of the genes coding for cyclins and cyclin-dependent kinases (Supplementary Figure S2B), centromeres (Supplementary Figure S2C) and core histone proteins (Supplementary Figure S2D). These findings suggest that loss of SMCHD1 reduces the expression of genes associated with cell cycle progression and may affect the expansion of myoblasts. To explore this possibility, we examined the expansion of SMCHD1-depleted myoblasts (Figure 2C-D). Measuring the rate of EdU incorporation into newly synthesized DNA, we observed that loss of SMCHD1 resulted in fewer myoblasts passing through S-phase of the cell cycle (Figure 2C). This decreased passage through cell cycle resulted in a reduced expansion of the progenitor population as determined by direct counting of cell numbers over time (Figure 2D). Taken together this data indicates that the loss of SMCHD1 in human myoblasts leads to a decreased ability of the progenitor population to expand their numbers.

**Figure 2.**
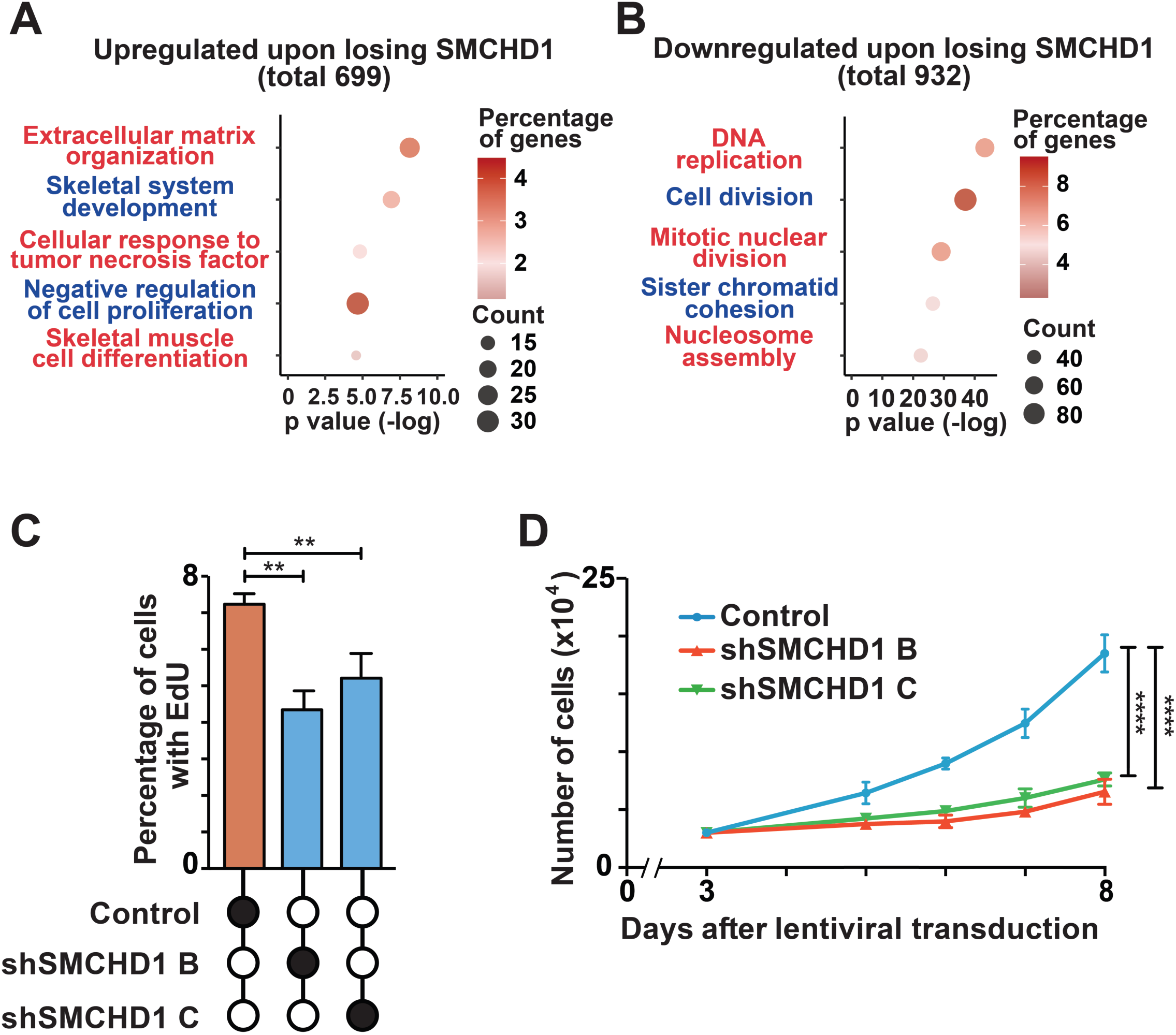
Loss of SMCHD1 in myoblasts leads to decrease in proliferation. **(A)** GO terms of up-regulated genes after SMCHD1 depletion. **(B)** GO terms of down-regulated genes after SMCHD1 depletion**. (C)** Five days after lentiviral transduction, EdU cell proliferation essay was performed and the percentage of EdU^+^ cells were quantified. Error bars represent standard deviation from 3 independent experiments. **(D)** An equal number of myoblasts was seeded 3 days after lentiviral transduction, and cell number was counted from day 5 to day 8 after lentiviral transduction. Error bars represent standard deviation from 3 independent experiments.

Having observed that loss of SMCHD1 results in FSHD-like changes in PAX7 target gene expression, we next examined whether the down-regulation of cell cycle genes observed upon depletion of SMCHD1 in myoblasts might be an unappreciated defect in myoblasts from FSHD patients with SMCHD1 mutations. Published datasets from FSHD patients with D4Z4 contractions (FSHD1–-28, 30) or SMCHD1 mutations (FSHD2–-31) were analyzed in combination with RNA-Seq data from our SMCHD1-depleted myoblast using Gene Set Enrichment Analysis (GSEA). Since these studies looked at myoblast differentiation, we limited our comparative analysis to the 0h time point of these published datasets (28, 30–31). Interestingly, we observed that genes that change their expression (both up- or down-regulated) in myoblasts upon loss of SMCHD1 overlapped significantly with genes that change their expression in FSHD myoblasts compared to healthy controls (Figure 3A and Supplementary Figure S3A). This strong correlation between FSHD patient myoblasts and myoblasts depleted of SMCHD1 suggested these cells may share similar defects, and led us to examine whether the expression of cell cycle genes was also dysregulated in FSHD patients. GO analysis showed that FSHD2 myoblasts from patients have altered expression of genes involved in cell division, and DNA replication (Supplementary Figure S3B), as well as negative regulation of cell proliferation (Supplementary Figure S3C). Unexpectedly, we observed that the changes in gene expression related to cell cycle and cell proliferation were specific to FSHD2 (Figure 3B), but not in FSHD1 myoblasts (Supplementary Figure S3D-F). The defect in cell cycle gene expression could also be observed (Figure 3C) in biopsies taken from FSHD2 patients (29, 32). Thus, down-regulation of genes involved in cell cycle progression is a characteristic shared between myoblasts depleted of SMCHD1 (myoblasts with shRNA-mediated depletion of SMCDH1 or FSHD2 patient samples), but not myoblasts from FSHD patients that retain SMCHD1 (FSHD1). The similar changes in cell cycle gene expression between SMCHD1-depleted myoblasts and in FSHD2 patient samples was intriguing given that previous studies have demonstrated that the doubling time of healthy and FSHD1 myoblasts were similar (33–34).

**Figure 3.**
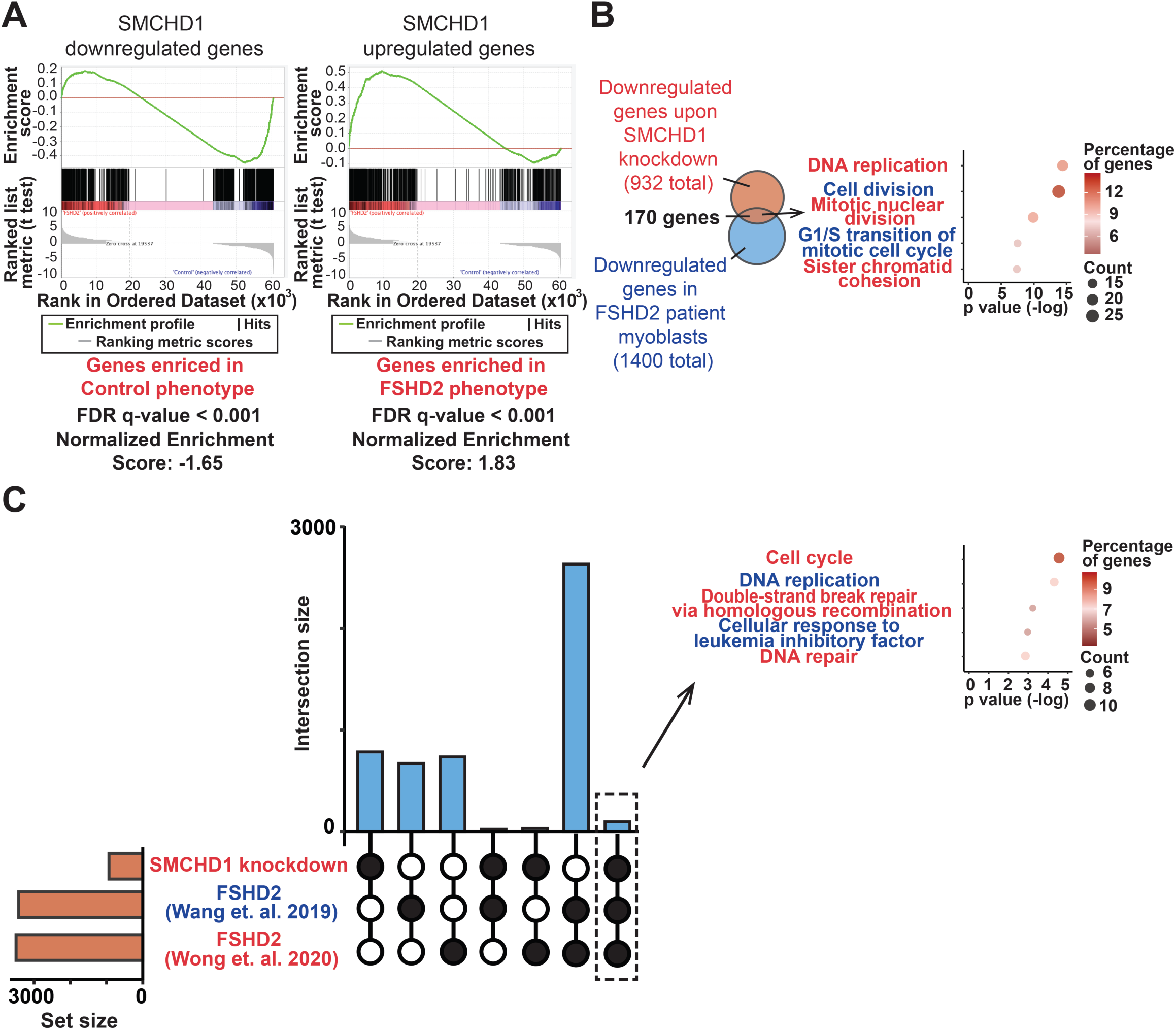
Comparison of changes in gene expression profiles upon SMCHD1 depletion and under FSHD2 condition. **(A)** Gene set enrichment analysis (GSEA) of DEGs in FSHD2 myoblasts compared to down-regulated (left panel) or up-regulated (right panel) genes upon SMCHD1 depletion. **(B)** (Left panel) Venn diagram of commonly down-regulated genes and (Right panel) dot plot of the top GO terms of the genes commonly down-regulated in SMCHD1-depleted myoblasts and in FSHD2 myoblasts. **(C)** (Left panel) Upset plot showing the commonly down-regulated genes upon SMCHD1 depletion and in FSHD2 biopsies from Wang et. al. (32) and Wong et. al. (29) studies. (Right panel) GO terms of commonly down-regulated genes upon SMCHD1 depletion and in FSHD2 biopsies.

### SMCHD1 binds at transcriptional regulatory regions of cell cycle genes in human myoblasts

SMCHD1 is known as a transcriptional silencer, yet the loss of SMDHD1 results in the down-regulation of cell cycle genes. This led us to examine whether genes down-regulated by the loss of SMCHD1 might be directly targeted by the epigenetic modifying protein. Probing the genome-wide binding of SMCHD1 using chromatin immunoprecipitation sequencing (ChIP-Seq), we identified 15560 sites (q value <= 0.05) of binding of SMCHD1 in proliferating myoblasts (Supplementary File S2). Knowing that SMCHD1 is involved in X chromosome inactivation (2), we first examined the distribution of SMCHD1 across the different chromosomes. As expected, we found that SMCHD1 binding was enriched on the X chromosome, with ∼4 fold more SMCHD1 binding sites relative to individual autosomes (15.1 vs 3.6 sites per million base pair) (Supplementary Figure S4A). Similarly, we observed a strong enrichment of SMCHD1 (Supplemental Figure 4B-4D) at sites previously known to be targeted by SMCHD1, including telomeres (35) as well as the 5S rRNA and tRNA clusters on chromosome 1 (36). This established that SMCHD1 in human myoblasts localizes to many of regions previously described for other cell types and provides confidence in the specificity of our ChIP-Seq.

We next wanted to associate SMCHD1 binding at specific genes to changes in gene expression in order to identify direct target genes. For this purpose, we performed Binding and Expression Target Analysis (BETA) which directly integrates data from ChIP-Seq and RNA-Seq (37). Using this approach, we identified 358 genes bound by SMCHD1 that changed their expression upon SMCHD1 depletion (Supplementary File S2). Though SMCHD1 has known silencing activity, we observed that many target genes are repressed upon loss of SMCHD1 (181 down-regulated and 177 up-regulated). Relating our SMCHD1 binding data to published datasets that had previously identified enrichment of histone modifications in human primary myoblasts (38), we observed that SMCHD1 preferentially bound at heterochromatic regions enriched for H3K9me3 marks (Figure 4A and Supplementary Table S4). This finding was expected and is consistent with the binding of SMCHD1 to the telomers and the inactive X chromosome, and work showing that the binding of SMCHD1 at the *D4Z4* locus is H3K9me3 dependent (39). Unexpectedly, we also observed an enrichment of SMCHD1 peaks at sites enriched for the transcriptionally permissive H3K4me3 mark (Figure 4A and Supplementary Table S4). This finding led us to examine whether SMCHD1 might be associated with transcriptional regulatory regions. Using the human GeneHancer database (40), our analysis determined that 38% of SMCHD1 direct target genes (Z score = 22.7) showed SMCHD1 binding at previously identified promoters or enhancers (Figure 4B). Consistent with previous findings (41) we observed the co-binding of CTCF at many of these SMCHD1-bound enhancers (Supplementary Figure S4E). This indicates a role for SMCHD1 in helping establish active enhancers at genes involved in cell cycle regulation.

**Figure 4.**
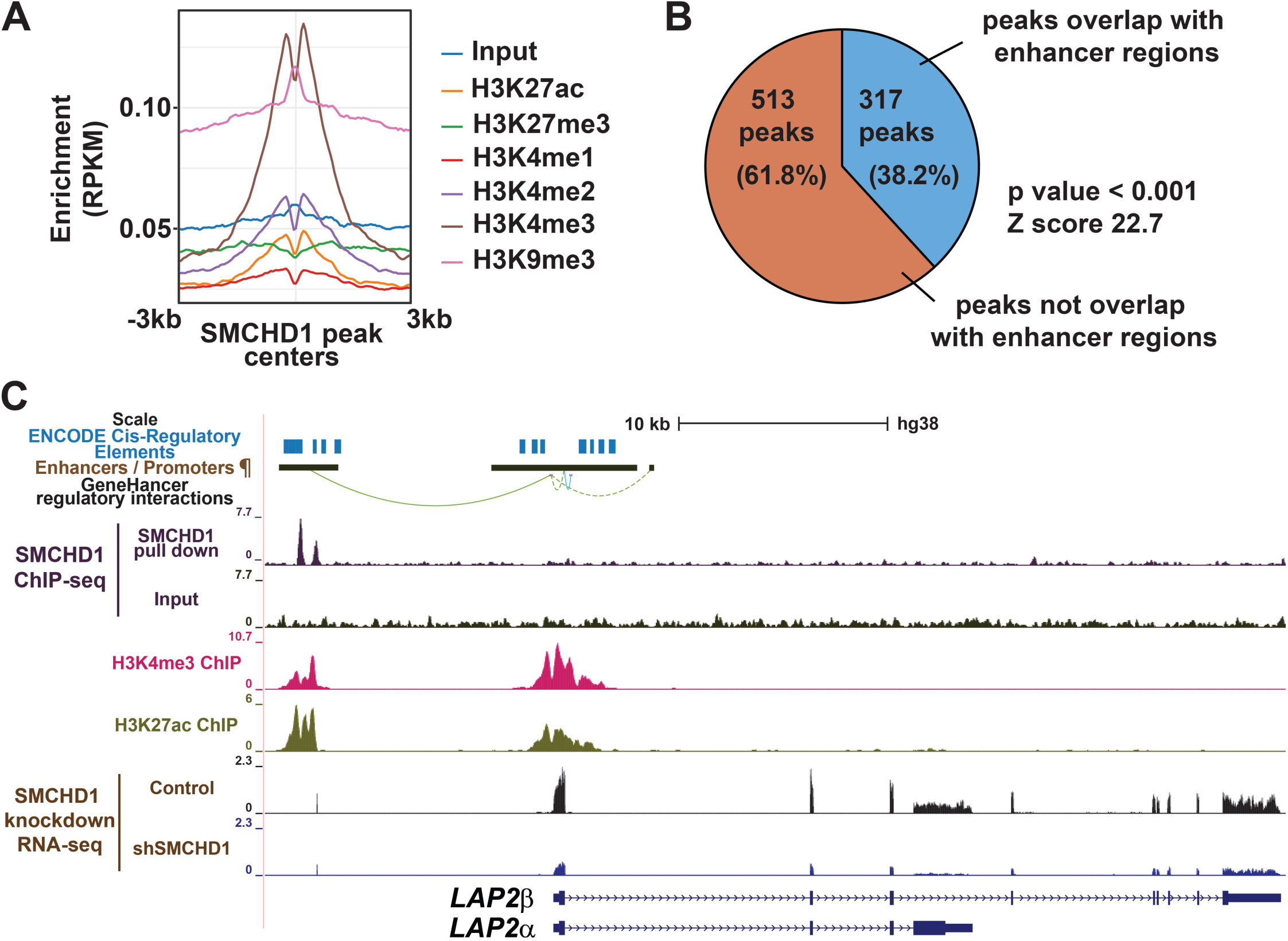
Binding regions of SMCHD1 in the human myoblast genome. (**A**) The enrichment of H3K9me3, H3K4me1, H3K4me2, H3K4me3, H3K27me3 and H3K27ac around SMCHD1 peak centers. (**B**) Percentage of direct targets-associated peaks that were located at promoters and enhancers. p value and Z score were calculated by permutation test using regioneR. **(C)** UCSC genome browser tracts showing ENCODE cis-regulatory elements, GeneHancer annotated enhancer / promoter regions (¶) and their regulatory interactions, as well as ChIP-Seq (SMCHD1, H3K4me3 and H3K27ac) and RNA-Seq reads (control: proliferating myoblasts expressing non-silencing scrambled shRNA, shSMCHD1: proliferating myoblasts expressing shRNA targeting *SMCHD1*) near the *LAP2* gene.

### SMCHD1 regulates proliferation of myoblasts via its direct target gene *LAP2*

Having identified a list of genes involved in cell cycle regulation as direct targets of SMCHD1, we next sought to identify genes that contribute to the cell proliferation defect in SMCHD1-depleted cells. Amongst these cell cycle regulators, we focused on Lamina-associated polypeptide 2 (LAP2, also known as Thymopoietin or TMPO) (Figure 4C), a direct target gene that is down-regulated in both SMCHD1-depleted myoblasts and FSHD2 patients (Figure 4C and 5A) while being unaltered in FSHD1 patients (Supplementary Figure S5A). Among the different isoforms of the *LAP2* gene (Figure 5B), LAP2α is a lamin A/C binding protein that has been shown to be essential for the proliferation of glioblastoma cells (42), while LAP2β is essential for nuclear envelope reassembly after mitosis in *xenopus* (43). We therefore tested whether the loss of LAP2 would also lead to a proliferation defect in human myoblasts. We were able to obtain a strong depletion of LAP2 expression in myoblasts using 2 different shRNA targeting LAP2 (Figure 5B, and Supplementary Figure S5B). Examining cell morphology at 9 days after shRNA treatment, we observed that myoblasts depleted of LAP2 became more flattened and spread out compared to control cells (expressing non-silencing shRNA) which continued to maintain a normal myoblast morphology (Figure 5C). Counting the cell numbers as the population proliferated over 10 days, myoblasts depleted in LAP2 showed a reduced rate of proliferation (Figure 5D). This indicates that the effect of losing SMCHD1 on the proliferation and morphology of myoblasts can be recapitulated by the loss of LAP2. Examination of exon inclusion from RNA-seq data indicated that LAP2β is the predominant isoform of the gene expressed in proliferating human myoblasts (Figure 4C). To determine whether LAP2 could rescue the SMCHD1 depletion-mediated proliferation defect, we overexpressed LAP2β in myoblasts depleted of SMCHD1. Excitingly, ectopic expression of LAP2β was sufficient to significantly improve the proliferation of SMCHD1-depleted myoblasts (Figure 5E). In summary, our data suggest that SMCHD1 helps ensure the efficient proliferation of human myoblasts by activating the expression of its target genes involved in cell cycle progression, including LAP2.

**Figure 5.**
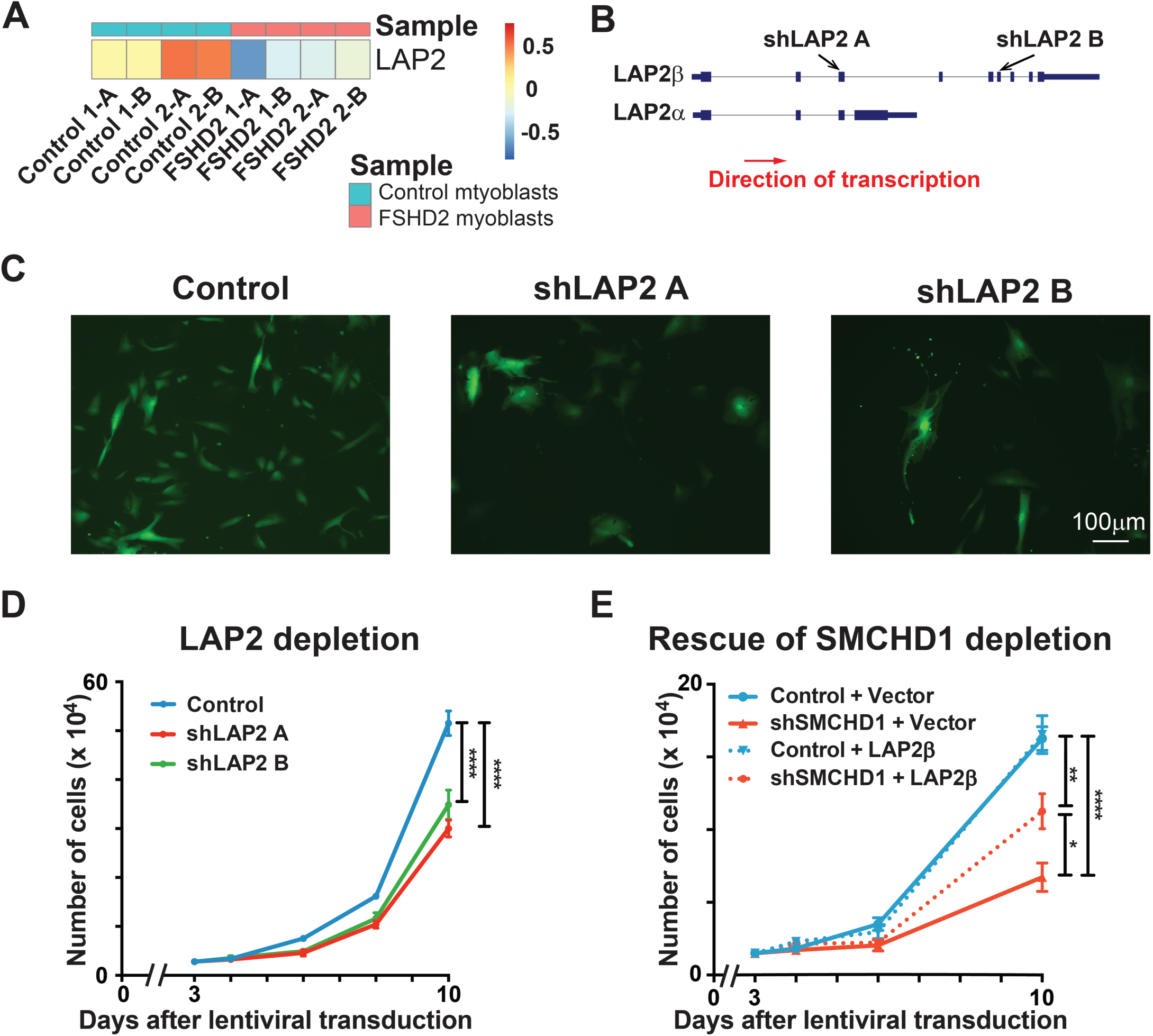
LAP2 depletion slows myoblast proliferation. **(A)** Heatmap showing the RNA expression of *LAP2* in FSHD2 and control non-FSHD myoblasts (GEO accession number GSE143493). **(B)** Different *LAP2* isoforms and target regions of different *LAP2* shRNAs. (**C**) Nine days after *LAP2* shRNA transduction, cells were observed under fluorescence microscope for GFP^+^ shRNA expressing cells. **(D)** Three days after *LAP2* depletion, equal number of myoblasts were seeded on 6 well plates. Cell number was counted on day 4, 6, 8 and 10 after lentiviral transduction. Error bars represent standard deviation from 3 independent experiments. **(E)** Lentivirus was added to the culture medium of proliferating myoblasts as indicated (control: non-silencing scrambled shRNA, shSMCHD1: shRNA targeting *SMCHD1*, vector: plenti-GIII vector alone, LAP2β: plenti-GIII vector expressing LAP2β transcript). Three days after lentiviral transduction, equal number of myoblasts were seeded on 6 well plates. Cell number was counted on day 4, 6 and 10 after lentiviral transduction. Error bars represent standard deviation from 3 independent experiments.

## DISCUSSION

The detrimental effects of reduced SMCHD1 levels in mature muscle myofibers have been shown to be at the heart of the FSHD disease pathology in FSHD2 patients. However, the importance of the ubiquitously expressed SMCHD1 protein in maintaining and expanding the myoblast population has yet to be fully established. Here we show that SMCHD1 is required for the efficient expansion of muscle progenitor cells. We found that myoblasts depleted of SMCHD1 show a reduced rate of proliferation that correlates with reduced expression of cell cycle-related genes at the transcription level. While these changes in gene expression occur in the absence of DUX4 expression, it is notable that the same dysregulation of cell cycle gene expression can be observed in muscle progenitor cells obtained from FSHD patients with SMCHD1 mutations. This is consistent with FSHD being a satellite cell-opathy (44–45) and suggests that impaired expansion of myoblasts during regeneration may contribute to the disease pathology in FSHD2 patients.

The application of a knockdown approach to diminish SMCHD1 levels has allowed us to delineate a role for the chromatin-modifying enzyme in regulating the proliferation of human muscle progenitor cells. The reduced expansion of myoblasts upon reduction of SMCHD1 levels is associated with the down-regulation of genes involved in DNA replication and cell cycle progression. Consistent with a role for SMCHD1 in promoting cell proliferation, studies in the early mouse embryo have shown that SMCHD1 is required for the expression of cell proliferation genes and that depletion of maternal SMCHD1 in the oocyte reduces the number of cells present in the blastocyst (46). Similarly, deletion of SMCHD1 in the hematopoietic system resulted in hematopoietic stem cells (HSCs) that were inefficient at competing with wild-type HSCs for repopulating the bone-marrow niche (47). This suggests that SMCHD1 plays a role in the expansion of various stem cell populations. We identified *LAP2* as a key SMCHD1-target gene that ensures efficient proliferation of the myoblasts. Indeed, SMCHD1 binds to a putative enhancer for *LAP2*, and is required for efficient expression of this protein that links the nuclear envelope to chromatin. The LAP proteins have been shown to play an important role in cell proliferation through the regulation of Rb/E2F1 activity as well as through the re-organization of nuclear envelope-associated proteins after cell division (48). Importantly, we demonstrated that exogenous expression of the LAP2β isoform was able to partially rescue the proliferation defect observed in conditions of diminished SMCHD1 levels. Our findings are particularly relevant for patients with FSHD2 since the diminished expression of *LAP2* was also observed in individuals carrying SMCHD1 mutations. Although it is generally accepted that FSHD2 patients do not present unique clinical phenotypes (49), we observed that muscle progenitor cells from FSHD2 patients show reduced expression of many of cell cycle-related genes that were identified upon SMCHD1 depletion. As such, patients with mutations in SMCHD1 are expected to have defects in their ability to repair the damaged myofibers in addition to the loss of muscle due to spurious DUX4 expression. Though patients with FSHD1 have been shown to have defective regeneration due to impaired muscle differentiation (45), this identification of this DUX4-independent role for SMCHD1 in muscle regeneration (Figure 6), provides an additional explanation for previous findings that SMCHD1 mutations can act as a disease modifier that increases the disease severity FSHD1 patients with D4Z4 locus contractions that cause muscle wasting (17). In these patients, not only would the muscle fibers undergo wasting but the muscle would also be defective in their ability to regenerate due to poor expansion of the muscle progenitors. Thus, we have identified a key role for SMCHD1 in maintaining the expression of genes involved in the proliferation of muscle progenitor cells.

**Figure 6.**
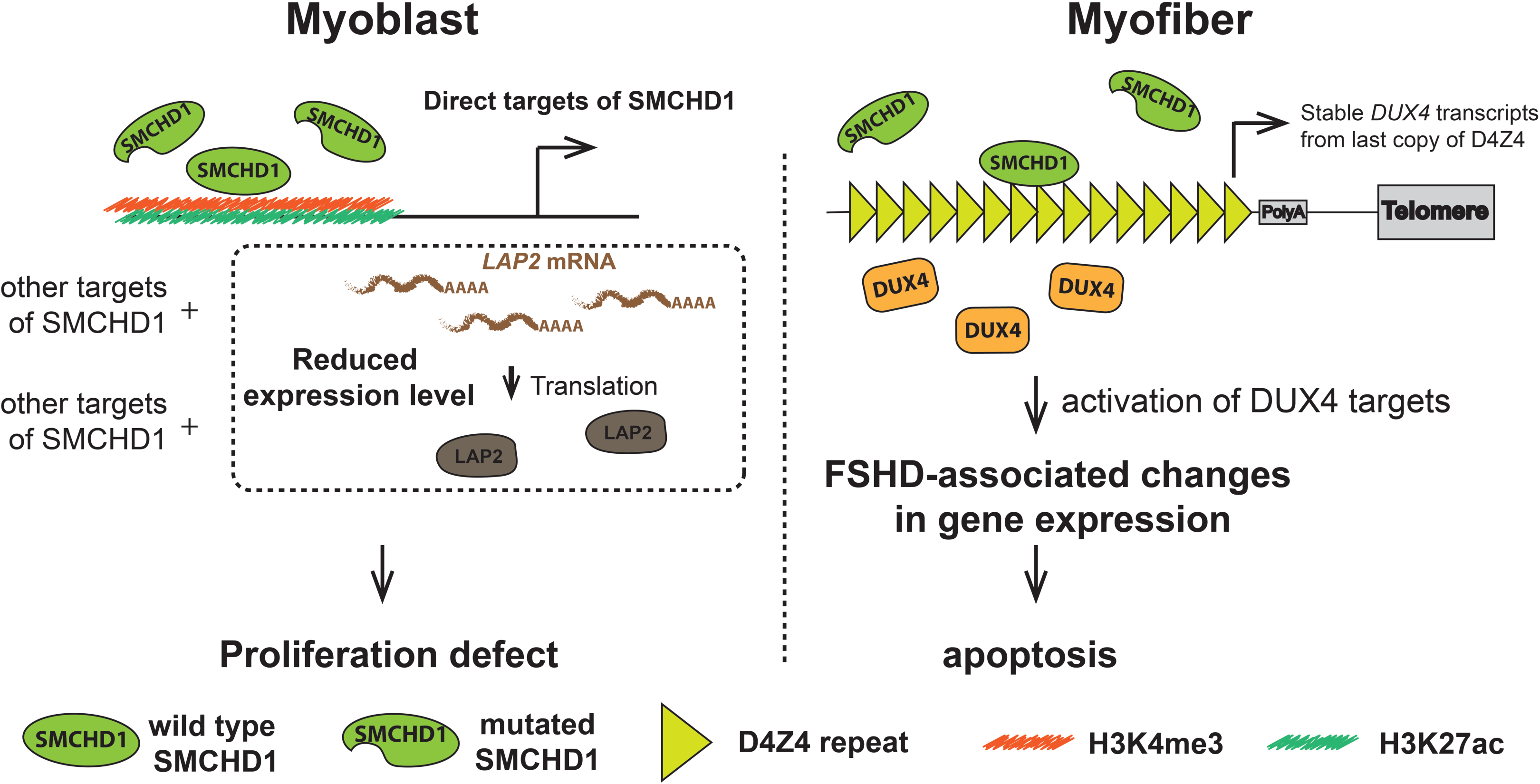
Working model of how the loss of functions of SMCHD1 result in FSHD2-associated defects. (Left panel) Loss of SMCHD1 binding on the promoter and enhancer regions of its target genes in myoblasts may result in additional problems to myoblasts growth, on top on those caused by spurious DUX4 expression. **(Right panel)** Heterozygous SMCHD1 mutations in FSHD2 patients lead to a reduced SMCHD1 binding on the D4Z4 locus and the expression of DUX4 in muscle, which results in the degeneration of myofiber.

A surprising finding from our study was the determination that SMCHD1 plays the role of a transcriptional activator in certain contexts. SMCHD1 is classically known for its role in silencing large chromatin domains (1). During the onset of X-chromosome inactivation, recruitment of SMCHD1 to the intermediate compartments along the chromosome allows their fusion to form repressive megadomains that ensure gene silencing (3). As a non-classical SMC protein, it is thought that SMCHD1 creates these repressive megadomains by entrapping DNA in dense loops of chromatin (50). However, we and others have observed that depletion of SMCHD1 results in a large number of genes being down-regulated, a result consistent with a transcriptional activator (this study; 46-47). As expected, genome-wide analysis of SMCHD1 binding showed an accumulation at sites-specific repressive domains in myoblasts. Unexpectedly, we also observed that SMCHD1 binds at transcriptional enhancers and/or promoters for many of the genes that show down-regulation upon removal of the epigenetic modifier. This suggests that SMCHD1 can act as a transcriptional repressor or activator depending on the genomic context. Of note, SMCHD1 is not the first example in which an epigenetic regulator has been found to have a dual role in regulating gene expression. Indeed, the polycomb repressive complexes PRC1 and PRC2 play an important role in maintaining transcriptional repression at the *D4Z4* locus (51–53). However, recent studies have shown that the binding of PRC1 (54–56) and PRC2 (57–58) at enhancers can also promote the activation of developmental gene expression by establishing long-range chromatin interactions. Interestingly, SMCHD1 is known to be targeted to sites along the X-chromosome through interaction with PRC1 and the resulting H2AK119ub marks the complex deposits (59–60). It is tempting to speculate that PRC1 may also be responsible for targeting SMCHD1 to specific enhancers for gene activation. Recruitment of SMCHD1 to these regions could then bring the promoter into close contact with the SMCHD1/PRC1-bound enhancer through a loop extrusion mechanism. This scenario would provide a molecular explanation of how PRC1 is able to bring together DNA elements to mediate long-range interactions. However, generating evidence for such a mechanism PRC1/SMCHD1-mediated gene activation will require further studies.

In summary, we have identified a role for SMCHD1 in modulating the expansion of muscle progenitor cells through the activation of genes that control cell cycle progression. These findings suggest that impaired muscle regeneration may contribute to the disease pathology in FSHD2 patients with mutations that induce SMCHD1 haploinsufficiency.

## ACKNOWLEDGEMENT

We thank Odile Neyret for ChIP-Seq and RNA-Seq library sequencing; Hina Bandukwala for help with TelomereHunter analysis, and Lifang Li for technical help. This work was funded by the ERA-Net for Research on Rare Diseases [D.G. and F.J.D.]; and the Canadian Institutes of Health Research [FDN-143330 to F.J.D.].

