## Supplemental Files for "SMCHD1 activates the expression of genes required for the expansion of human myoblasts"

**This PDF file includes:**

Materials and Methods
Figures S1 to S5
Supplemental Tables S1-S4
References 1-21

**RT-qPCR**

Total RNA from cultured cells was harvested using RNA-STAT60 reagent (Tel-Test # CS-111) according to the manufacturer's protocol. Purified RNA (2000 ng) was subjected to reverse transcriptase reaction in the presence of 2.5 mM dNTP (Thermo Fisher Scientific #10297-018), 30 ng/ml random primers (Thermo Fisher Scientific # 48190011) with 6 U/µl Moloney Murine Leukemia Virus Reverse Transcriptase (New England BioLabs # MO253L). qPCR reactions were performed using PowerUP SYBR Green Master Mix (Thermo Fisher Scientific # A25778) according to the user manual. Expression of genes were normalized to GAPDH expression level using the delta-delta Ct method. *p<0.05, **p<0.01, ***p<0.001, ****p<0.0001. Statistical quantification was performed using student's two-tailed t-test on the statistical graphing software GraphPad Prism v6.0.

Primers used in qPCR:

| **Primer name** | **Sequence** |
|---|---|
| SMCHD1-for | CGACAGATTGTCCAGTTCCTC |
| SMCHD1-rev | CCAATGGCCTCTTCTCTCTG |
| GAPDH-for | TCAAGAAGGTGGTGAAGCAGG |
| GAPDH-rev | ACCAGGAAATGAGCTTGACAAA |
| LAP2-for | CGGACTTCTCCAGTGACGA |
| LAP2-rev | GGACCAGGATTCACTCCGTA |
| CCNA2-for | CTCTACACAGTCACGGGACAAAG |
| CCNA2-rev | CTGTGGTGCTTTGAGGTAGGTC |
| CCNB1-for | GACCTGTGTCAGGCTTTCTCTG |
| CCNB1-rev | GGTATTTTGGTCTGACTGCTTGC |
| CCNB2-for | CAACCAGAGCAGCACAAGTAGC |
| CCNB2-rev | GGAGCCAACTTTTCCATCTGTAC |
| CDK1-for | GGAAACCAGGAAGCCTAGCATC |
| CDK1-rev | GGATGATTCAGTGCCATTTTGCC |
| CDKN3-for | ATGGAGGGACTCCTGACATAGC |
| CDKN3-rev | TCTCCCAAGTCCTCCATAGCAG |
| CENPA-for | GGCGGAGACAAGGTTGGCTAAA |
| CENPA-rev | GGCTTGCCAATTGAAGTCCACAC |
| CENPI-for | GCCTTTGTTGTCGGTCACAGTTC |
| CENPI-rev | AGAAGAGCTGCGCTAGATGGTC |
| CENPE-for | GGAGAAAGATGACCTACAGAGGC |
| CENPE-rev | AGTTCCTCTTCAGTTTCCAGGTG |
| HIST1H3A-for | TCCGCCGTTATCAGAAGTCCAC |
| HIST1H3A-rev | GCTCTGGAAACGCAGGTCTGTT |
| HIST1H2AB-for | GGCGGTGCTTGAGTACCTGAC |
| HIST1H2AB-rev | AAGCTCCTCGTCATTGCGGATG |
| HIST1H1B-for | CCGAAAAAGGCAACCAAGAGTCC |
| HIST1H1B-rev | GTTTTCACACGCCAGCTTCCTAC |
| HIST1H3B-for | GCGAGAAATCGCCCAAGACTTC |
| HIST1H3B-rev | CAAAGAGCCCTACCAAGTAGGC |

**Immunoblotting**

Cells were lysed in RIPA buffer (150 mM NaCl, 1% NP-40, 0.5% sodium deoxycholate, 0.1% SDS. 50 mM Tris, pH 8.0) with 1x EDTA-free Protease Inhibitor Cocktail (Roche # 1187358000), left on ice for

5 minutes followed by centrifugation at 16,000 x g for 10 minutes at 4°C. Supernatants were collected and proteins were quantified using protein assay dye reagent (Biorad #5000006). Western blotting was performed as previously described (1), with 1:1000 SMCHD1 antibody (Bethyl Laboratories #A302-871A) and 1:000 α-Tubulin antibody (Cell Signaling Technology #3873) used in the primary antibody incubation step.

### Supplementary figures

#### Supplementary Figure S1

**(A)** Quantitative PCR results showing the relative expression of *SMCHD1* in myoblasts 5 days after transduction of lentiviral vectors expressing non-silencing shRNA (Control) or shRNA targeting *SMCHD1* (shSMCHD1 A, shSMCHD1 B and shSMCHD1 C). Error bars represent standard deviations of 3 independent experiments. **(B)** (Upper panel) Western blot showing the expression of SMCHD1 and (lower panel) α-tubulin of the same blot 5 days after lentiviral transduction. **(C)** DUX4 scores of samples transduced with non-silencing scrambled shRNA (Control) or *SMCHD1* targeting shRNA (shSMCHD1), based on DUX4 targets identified by Yao et. al. (2) and Geng et. al. (3). **(D)** Venn diagram showing the target genes repressed by PAX7 and differentially expressed upon SMCHD1 depletion. **(E)** Number of genes repressed by PAX7 that were down-regulated (red) and up-regulated (blue) upon SMCHD1 depletion.

#### Supplementary Figure S2

**(A)** Heatmaps showing the changes in expression of genes encoding collagen subtypes upon losing SMCHD1. **(B-D)** Quantitative PCR results showing the relative expression of mRNA code for (B) cyclins and cyclin dependent kinases, (C) centromere proteins and (D) core histones 5 days after transduction of lentiviral vectors expressing non-silencing scrambled shRNA (Control) or shRNA targeting SMCHD1 (shSMCHD1 A, shSMCHD1 B and shSMCHD1 C). Error bars represent standard deviations of 3 independent experiments.

#### Supplementary Figure S3

**(A)** Gene set enrichment analysis (GSEA) of DEGs in FSHD1 myoblasts compared to down-regulated (left panel) or up-regulated (right panel) genes upon SMCHD1 depletion. **(B-C)** Dot plots showing the top GO terms of (B) down-regulated and (C) up-regulated genes in primary FSHD2 myoblasts (differentiation day 0) RNA-Seq dataset (GEO accession number GSE143493). **(D)** (Left panel) Venn diagram of commonly down-regulated genes and (Right panel) dot plot of top GO terms of the genes commonly down-regulated in SMCHD1-depleted myoblasts and in FSHD1 myoblasts. **(E-F)** Dot plots showing the top GO terms of (E) down-regulated and (F) up-regulated genes in FSHD1 myoblasts clones having shortened D4Z4 locus (GEO accession number GSE102812).

#### Supplementary Figure S4

**(A)** Average number of SMCHD1 peaks on autosomes and on chromosome X. **(B)** Telomere content in the input and SMCHD1 pull-down sample calculated by the TelomereHunter software. **(C-D)** UCSC genome browser tracks showing SMCHD1 ChIP-Seq reads and SMCHD1 depletion RNA-Seq reads (control: proliferating myoblasts expressing non-silencing scrambled shRNA, shSMCHD1: proliferating myoblasts expressing shRNA targeting *SMCHD1*) near (C) 5S rRNA cluster, (D) tRNA cluster. **(E)** (Upper panel) CTCF motif was enriched among the SMCHD1 peaks associated with direct targets. (Lower panel) Number of CTCF peaks colocalized with SMCHD1 peaks. p value and Z score were calculated by permutation test using regioneR.

#### Supplementary Figure S5

**(A)** Heatmap showing the expression of LAP2 from control non-FSHD and FSHD1 myoblasts (GEO assession number GSE102812). **(B)** Quantitative PCR results showing the relative expression of *LAP2* after transduction of lentiviral vectors expressing non-silencing shRNA (Control) or shRNA targeting *LAP2* (shLAP2 A and shLAP2 B). Error bars represent standard deviation from 3 independent experiments.

**Supplementary Table S1. List of reagents**

| Name of Regent | company name | Catalog # |
|---|---|---|
| α-Tubulin antibody | Cell Signaling Technology | 3873 |
| BGS | HyClone | SH30541.03 |
| DAPI | Sigma | D9542 |
| Dexamethasone | Sigma | D2915 |
| DMEM | HyClone | SH3024301 |
| dNTP | Thermo Fisher Scientific | 10297-018 |
| EDTA | Fisher Scientific | S311-500 |
| EDTA-free Protease Inhibitor Cocktail | Roche | 1187358000 |
| EdU Cell Proliferation Kit for Imaging | Thermo Fisher Scientific | C10340 |
| FGF | Stemcell Technologies | 78003.2 |
| Formaldehyde | Fisher Scientific | BPBP53125 |
| Gelatin | Sigma | G9391 |
| Glycerol | Fisher Scientific | G33-1 |
| Goat anti-rabbit IgG antibody conjugated to Alexa Fluor 488 | Thermo Fisher Scientific | A-11034 |
| HEPES | Sigma | H3784 |
| HGF | Stemcell Technologies | 78019.2 |
| IRDye 680RD donkey anti-Rabbit IgG antibody | Li-Cor | LIC-926-68073 |
| IRDye 800CW goat anti-Mouse IgG antibody | Li-Cor | LIC-926-32210 |
| KAPA HyperPrep Kit | Roche | 07962363001 |
| KAPA stranded RNA-seq kit | Roche | 07962169001 |
| M-MuLV Reverse Transcriptase | New England BioLabs | MO253L |
| Medium 199 | Thermo Fisher Scientific | 31100035 |
| Na-deoxycholate | Sigma | D6750 |
| NaCl | Sigma | S9888 |
| Normal rabbit IgG | Thermo Fisher Scientific | 10500C |
| NP-40 | Sigma | 74385 |
| PBS | Gibco | 21600-069 |
| pDONR223-LAP2β plasmid | Genome editing and molecular biology facility of the University of Ottawa | N/A |
| Penicillin/streptomycin | Wisent Bioproducts | 450-201-EL |
| pLenti-EF1a-Blank vector | Applied Biological Materials | LV588 |
| pMD2.G vector | Addgene | 12259 |
| Polybrene | Sigma | H9268 |

| | | |
|---|---|---|
| Polyethylenimine | Polysciences | 23966 |
| PowerUP SYBR Green Master Mix | Thermo Fisher Scientific | A25778 |
| Protein Assay Dye Reagent | Bio-Rad | 5000006 |
| psPAX2 vector | Addgene | 12260 |
| PureLink RNA mini kit | Thermo Fisher Scientific | 12183018A |
| Random Primers | Thermo Fisher Scientific | 48190011 |
| Ribo-Zero Magnetic Gold Kit | Illumina | MRZG12324 |
| RNA-STAT60 reagent | Tel-Test | CS-111 |
| SDS | Sigma | L3771 |
| SMCHD1 antibody (for ChIP-Seq) | Abcam | ab31865 |
| SMCHD1 antibody (for western blot) | Bethyl Laboratories | A302-871A |
| Sucrose | Fisher Scientific | S6500 |
| Tris base | Fisher Scientific | BP154-1 |
| Triton X-100 | Sigma | T8787 |
| Vitamin B12 | Sigma | V2876 |
| Zinc sulfate | Fisher Scientific | Z68500 |

**Supplementary Table S2. Software and databases used in data analysis**

| Software and algorithms | Reference | Web Site |
|---|---|---|
| BEDTools | (4) | https://bedtools.readthedocs.io/en/latest/ |
| BETA algorithm | (5) | http://cistrome.org/BETA/ |
| bowtie2 (v2.3.5.1) | (6) | http://bowtie-bio.sourceforge.net/bowtie2/index.shtml |
| DAVID (v6.8) | (7-8) | https://david.ncifcrf.gov |
| DESeq2 (v1.30.0) | (9) | https://bioconductor.org/packages/release/bioc/html/DESeq2.html |
| EnhancedVolcano (v1.6.0) | | https://bioconductor.org/packages/release/bioc/html/EnhancedVolcano.html |
| EnrichedHeatmap (v1.18.2) | (10) | https://bioconductor.org/packages/release/bioc/html/EnrichedHeatmap.html |
| Fiji (v2.1.0) | (11) | https://imagej.net/software/fiji/ |
| ggplot2 (v3.3.0) | (12) | https://ggplot2.tidyverse.org |
| GSEA software | (13-14) | https://www.gsea-msigdb.org/gsea/index.jsp |
| HOMER (v4.11) | (15) | http://homer.ucsd.edu/homer/motif/ |
| MACS2 | (16) | https://github.com/macs3-project/MACS |
| pheatmap (v1.0.12) | | https://cran.r-project.org/web/packages/pheatmap/index.html |

| | | |
|---|---|---|
| GraphPad Prism (v6.0) | | https://www.graphpad.com/scientific-software/prism/ |
| R (v4.0.0) | | https://www.r-project.org |
| regioneR (v 3.14) | (17) | http://bioconductor.org/packages/release/ bioc/html/regioneR.html |
| Rsubread (v1.6.2) | (18) | https://bioconductor.org/packages/release/ bioc/html/Rsubread.html |
| samtools | (19) | http://samtools.sourceforge.net |
| STAR (v2.7.5a) | (20) | https://github.com/alexdobin/STAR |
| TelomereHunter | (21) | https://pypi.org/project/telomerehunter/ |

**Supplementary Table S3. List of shRNA vectors**

The following shRNA vectors were purchased from GeneCopoeia (MD, USA):

| **shRNA Vector** | **Target sequence** |
|---|---|
| psi-LVRU6MH-shControl | GCTTCGCGCCGTAGTCTTA |
| psi-LVRU6MH-shSMCHD1 A | GGACGGTGTACTTGTTTGATC |
| psi-LVRU6MH-shSMCHD1 B | GGGATTATCCGTTATCATCCA |
| psi-LVRU6MH-shSMCHD1 C | CCTATTGGTGCGTTAAGAATT |
| psi-LVRU6GP-shControl | GCTTCGCGCCGTAGTCTTA |
| psi-LVRU6GP-shLAP2 A | GGAACAGAATCAAGATCTTCT |
| psi-LVRU6GP shLAP2 B | GCTGAAACTATAATGGCTTCA |

**Supplementary Table S4.**

Percentage of SMCHD1 peaks overlapped with histone modification marks. Overlapping were done using BedTools. p value and Z score were calculated by permutation test using regioneR.

| | **Number of peaks** | **Percentage of overlap** | **p value** | **Z-score** |
|---|---|---|---|---|
| Overlap with H3K9me3 | 5558/15560 | 35.72 | <0.001 | 230.69 |
| Overlap with H3K27me3 | 90/15560 | 0.58 | <0.001 | -8.87 |
| Overlap with H3K4me1 | 113/15560 | 0.73 | < 0.001 | -19.51 |
| Overlap with H3K4me2 | 768/15560 | 4.94 | < 0.001 | 12.48 |
| Overlap with H3K4me3 | 1167/15560 | 7.50 | < 0.001 | 54.95 |
| Overlap with H3K27ac | 414/15560 | 2.66 | 0.38 | -0.34 |

Supplementary figure S1

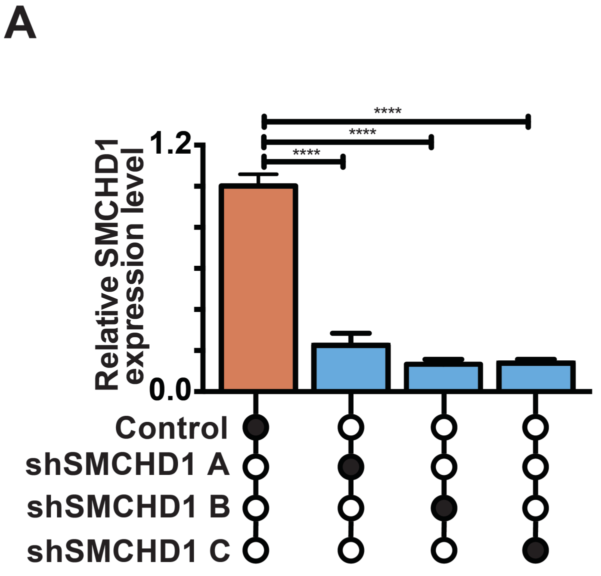


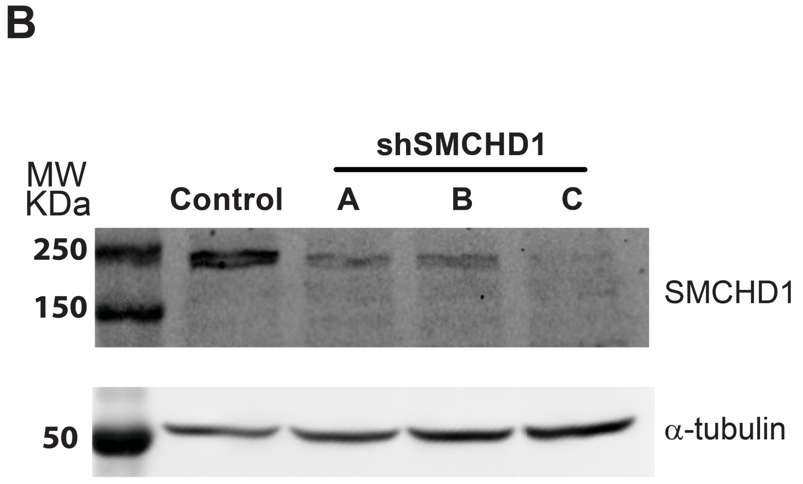


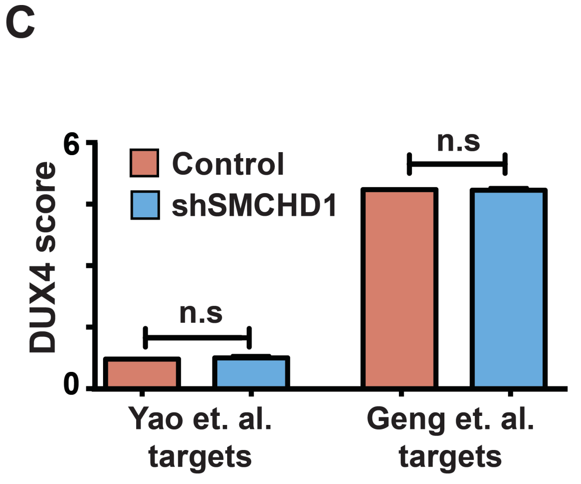


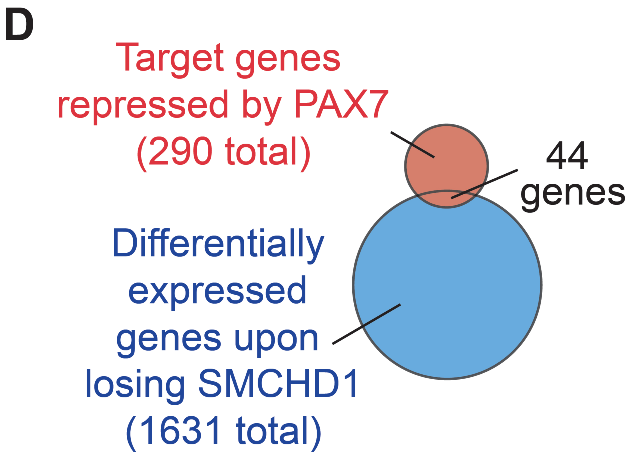


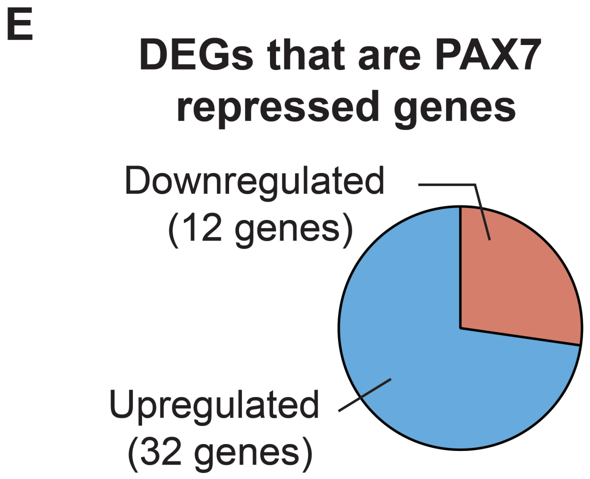

Supplementary figure S2

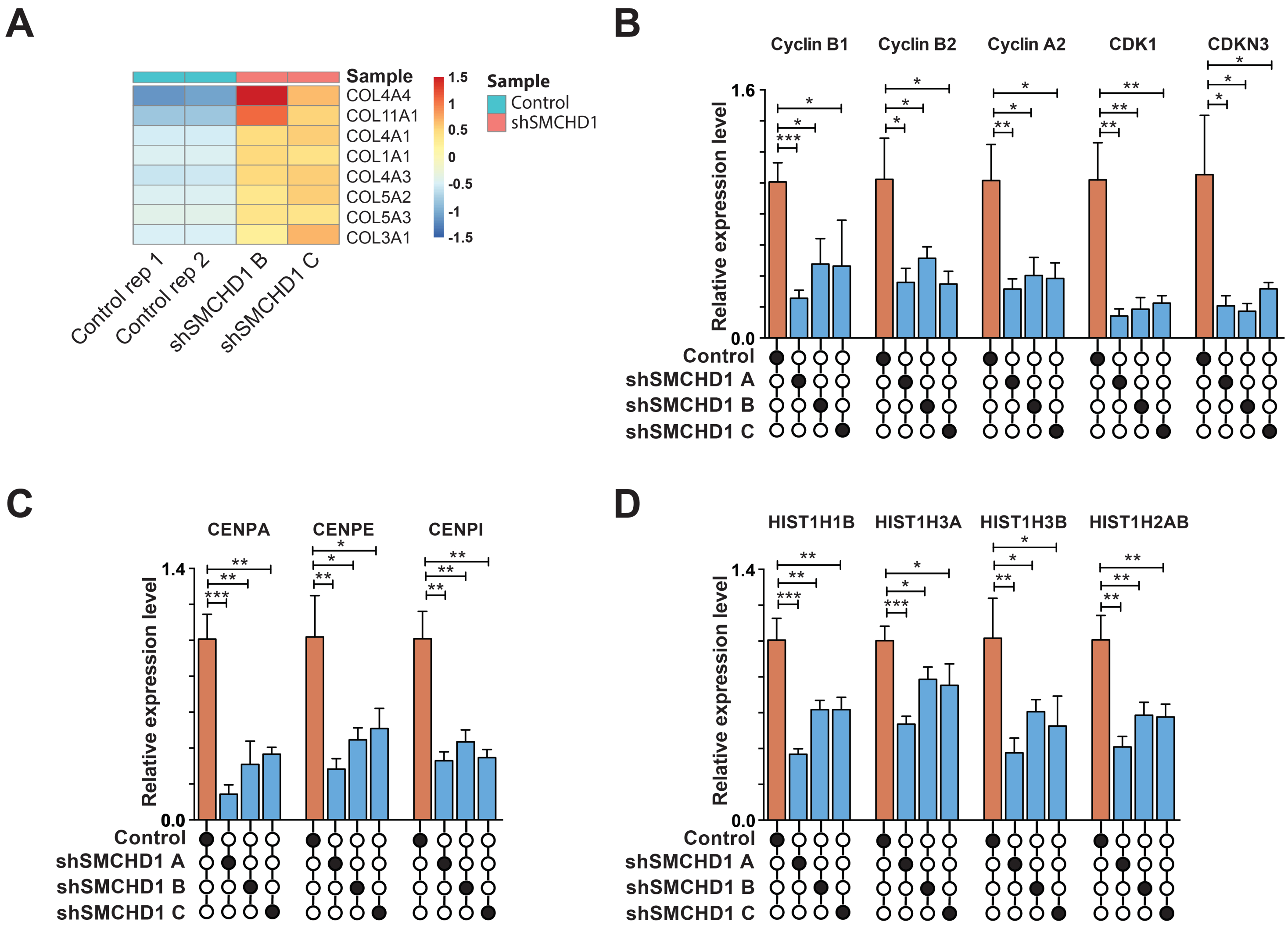

Supplementary figure S3

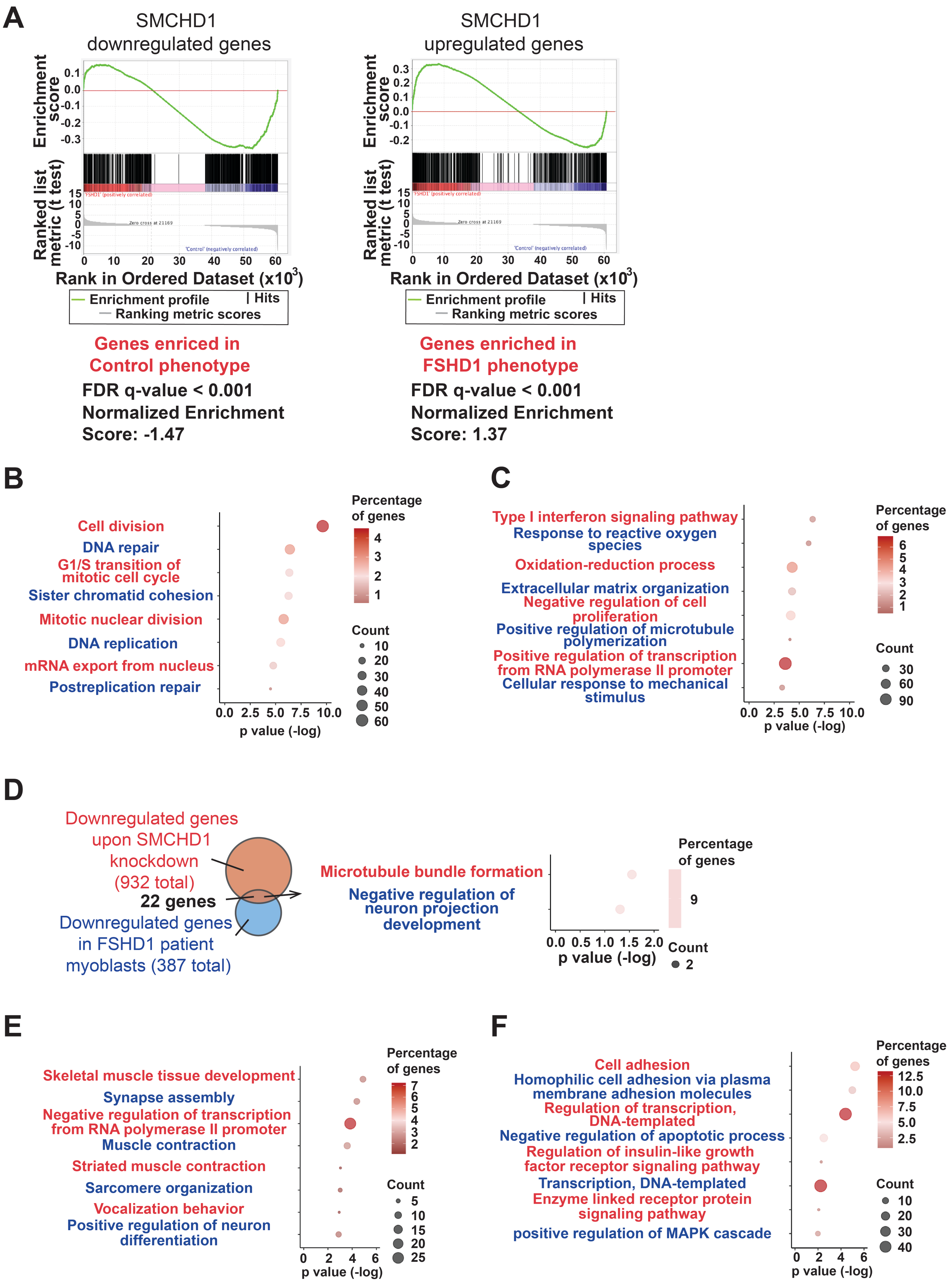

Supplementary figure S4

A

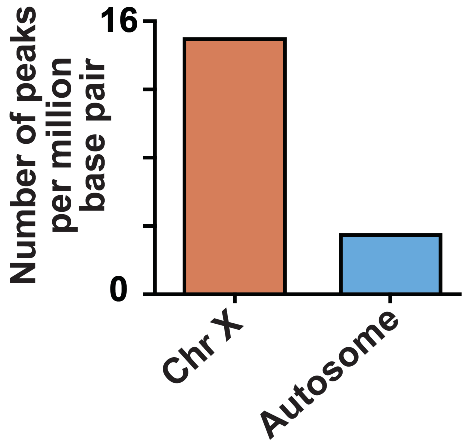


B

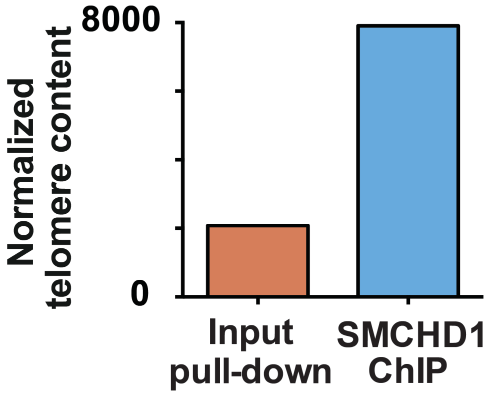


C

### 5S rRNA cluster

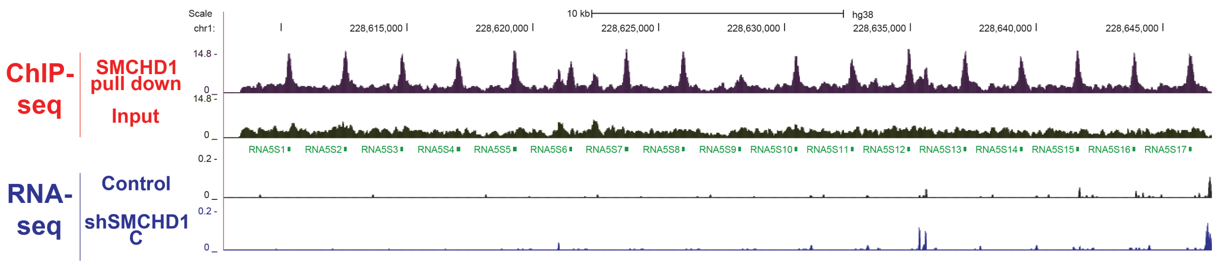


D

### tRNA cluster

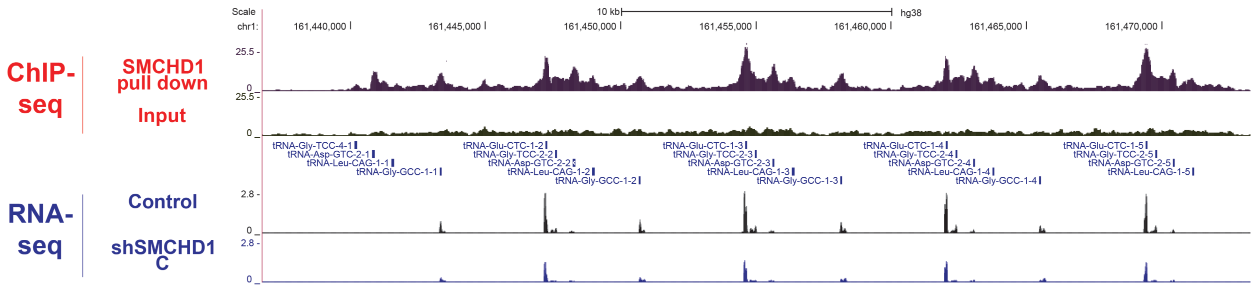


E

### Enriched motif in peaks associated with direct targets

| Motif | Name | p-value (Target VS non-target) | q-value (Target VS non-target) | % of target sequence with motif | % of non-target sequence with motif | % of GC normalized scramble sequence with motif |
|---|---|---|---|---|---|---|
| 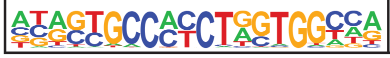 | CTCF | 1e-4 | 0.0108 | 7.35% | 4.07% | 0.36% |

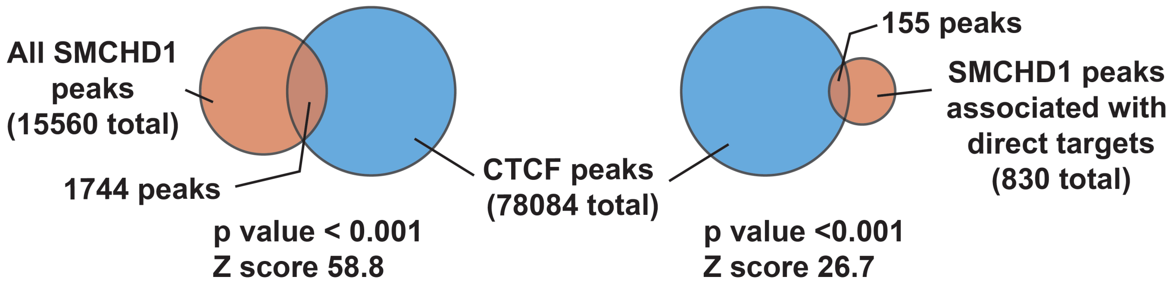

Supplementary figure S5

A

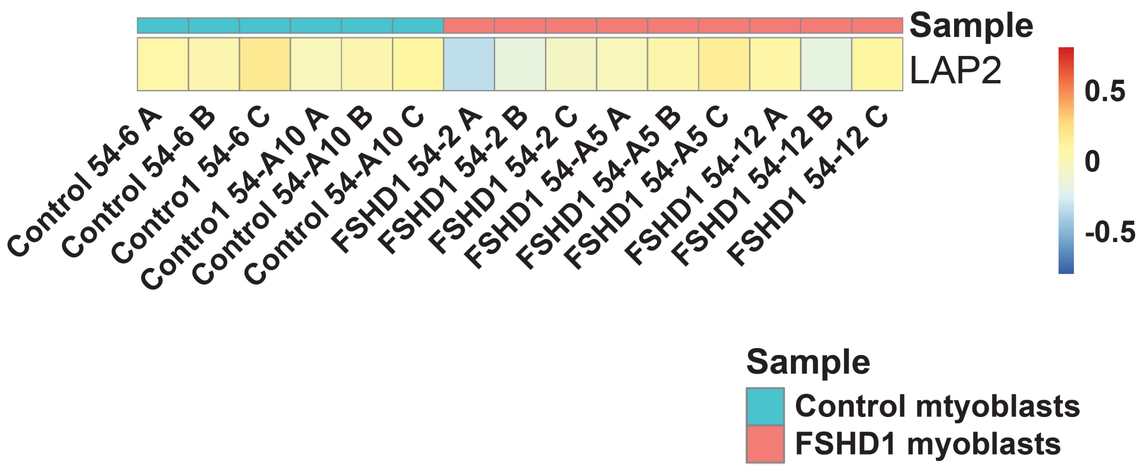


B

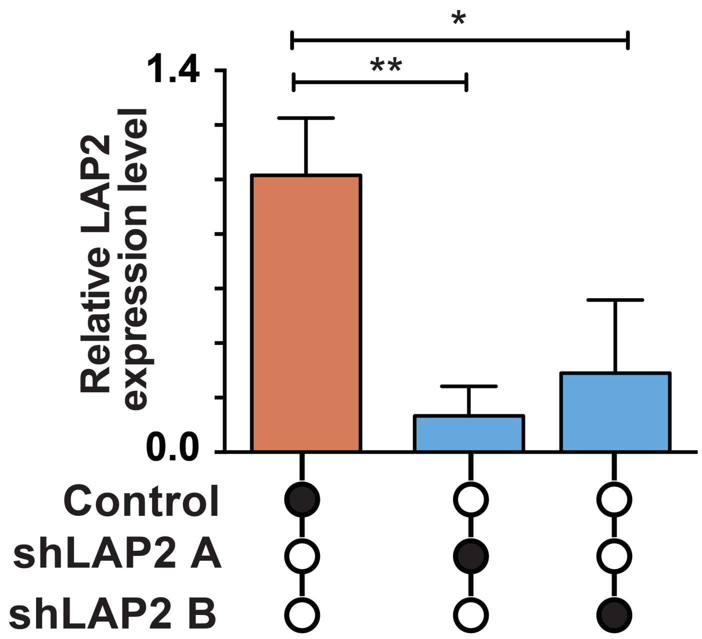
